# Retinal waves but not visual experience are required for development of retinal direction selectivity maps

**DOI:** 10.1101/2021.03.25.437067

**Authors:** Alexandre Tiriac, Karina Bistrong, Marla B. Feller

## Abstract

Retinal waves and visual experience have been implicated in the formation of retinotopic and eye-specific maps throughout the visual system, but whether either play a role in the development of the maps within the retina itself is unknown. We explore this question using direction-selective retinal ganglion cells, which are organized into a map that aligns to the body and gravitational axes of optic flow. Using two-photon population calcium imaging, we find that the direction selectivity map is present at eye opening and is unaltered by dark-rearing. Remarkably, the horizontal component of the direction selectivity map is absent in mice lacking normal retinal waves, whereas the vertical component remains normal. These results indicate that intrinsic patterns of activity, rather than extrinsic motion signals are critical for the establishment of direction selectivity maps in the retina.

**One Sentence Summary:** Horizontal direction selectivity in the retina is absent in mice lacking normal retinal waves.

## MAIN TEXT

Detecting the direction of visual motion, whether generated by self-motion or by objects moving within the visual field, is critical for everyday behavior. However, not all directions of motion are equally represented in the visual system. Several studies support the idea that the presence of directional visual stimulation instructs development of direction selectivity in visual cortex: horizontal motion detection develops after eye opening when animals become mobile (Jenks and Shepherd, 2020), visual deprivation leads to fewer direction-selective cells (Li et al., 2006), and training for particular directional motion at particular ages increases the proportion of direction-selective cells (Li et al., 2008; Roy et al., 2020).

In many species, direction selectivity is first manifested in the retina (Baden et al., 2020). In the mouse retina, direction-selective retinal ganglion cells (DSGCs) fire more action potentials in response to visual stimuli moving in one direction, called the preferred direction, than visual stimuli moving in the opposite direction, called the null direction (Barlow and Levick, 1965). In adult mice, the preferred directions of DSGCs cluster in four groups along two axes. The relative orientations of these axes vary with retinal location––following the axes of optic flow. Whereas the preferred directions of nasal- and temporal-preferring DSGCs follow the body axis, the preferred directions of dorsal- and ventral-preferring DSGCs follow the gravitational axis (Sabbah et al., 2017).

Given that the organization of retinal direction selectivity maps follow the axes of optic flow, does visual experience influence the establishment of this map? Previous work indicates that the direction-selective tuning of individual DSGCs is present at eye opening (Bos et al., 2016; Chan and Chiao, 2013; Elstrott et al., 2008; Wei et al., 2011; Yonehara et al., 2011) and is not diminished by dark-rearing (Bos et al., 2016; Chan and Chiao, 2008). In contrast, the impact of visual experience on the organization of direction selectivity maps is controversial. Work from our lab and others have described that dark-reared animals exhibit a significant difference in the clustering of preferred directions around cardinal axes defined by motion in visual field. (Bos et al., 2016; Chan and Chiao, 2013). However, these data were collected prior to the discovery that local direction selectivity maps change with retinal locations––following optic flow-generated axes (Sabbah et al., 2017). Therefore, direction selectivity map characterization requires careful account of retinal location, which may not have been considered in previous reports.

To assess the organization of direction selectivity maps across development, we performed two-photon calcium imaging over large areas of ventronasal and ventrotemporal retina (Figure 1A). We functionally classified DSGCs depending on whether they responded to both the onset and offset of the light (ON-OFF) or just to the onset of the light (ON; Figure 1B). ON-OFF and ON direction selectivity maps looks very similar, with the exception that ON maps have negligible numbers of nasal-preferring DSGCs (Figure 1C). In a subset of experiments, we used transgenic mice where ventral-preferring (Hb9-GFP) or nasal-preferring (Drd4-GFP) DSGCs were labelled with GFP.

**Figure 1.**
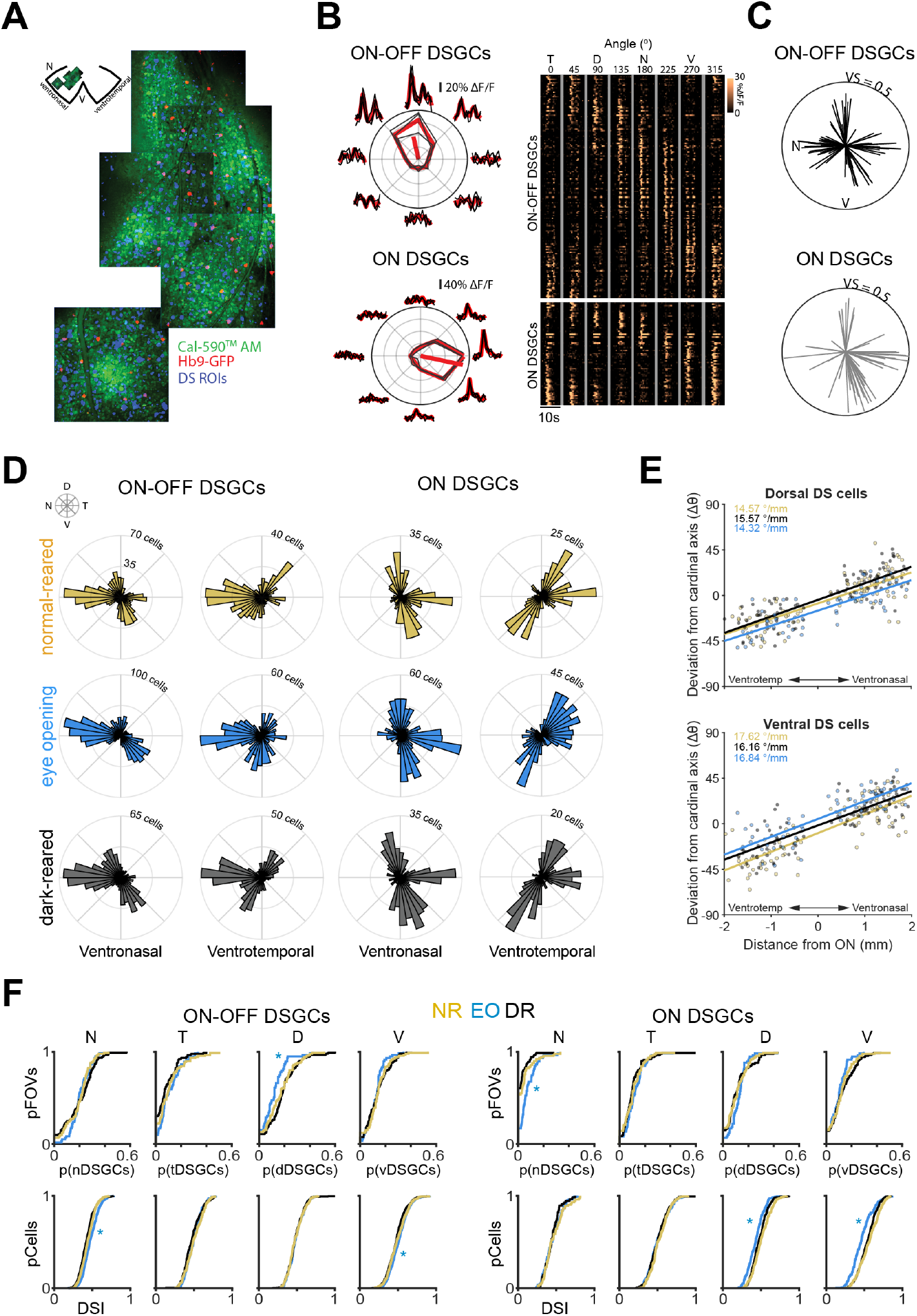
The direction selectivity map develops independent of visual experience. **A**. Schematic showing mapping of direction selectivity (DS) across a quadrant of the retina. As field of views are imaged, their locations are recorded and can be reconstructed post-hoc. **B**. Left: Tuning curve of an example ON-OFF and ON DSGC. Right: Heatmap of average responses to moving bars of ON-OFF and ON DSGCs identified from all of the FOVs in panel A. Each row is a different cell and are sorted based on preferred direction. **C**. Local DS map of the analyzed field of view shown in A for ON-OFF DSGCs (black) and ON DSGCs (grey). For each DS map, each vector is a single DSGC, with its direction representing the cell’s preferred direction and the length representing the strength of tuning. **D**. Polar histograms depicting the DS maps of normal-reared adults (NR, yellow; n = 2041 DSGCs across 10 mice), eye opening (EO, blue; n = 2472 DSGCs across 9 mice), and dark-reared adults (DR, black; n = 2261 DSGCs across 8 mice). The maps are segregated based on retinal location (ventronasal and ventrotemporal) and major class of DSGC (ON-OFF and ON). **E**. Scatter plots showing the rate of change of a functional group’s preferred direction (top: dorsal; bot: ventral) as a function of distance from the optic nerve. The slopes across the three groups are not statistically-significantly different. **F**. Top: Cumulative distribution plots depicting the different proportions of functional subgroups per field of view (N: nasal, T: temporal, D: dorsal, V: ventral) for ON-OFF and ON DSGCs and across different experimental groups. Bottom: cumulative distribution plots depicting the proportion of cells that exhibit different various DSI. * different from NR adult, p < 0.01.

In contrast to previous studies (Bos et al., 2016; Chan and Chiao, 2013), we find that direction selectivity maps are well-established by eye opening and do not degrade after dark-rearing (Figure 1D). To compare direction selectivity maps across conditions, we functionally clustered ON-OFF DSGCs and ON-DSGCs into 4 groups according to their preferred direction (see Methods, Figures S1, and Sabbah et al, 2017). In all three experimental groups, the preferred directions of vertical-preferring DSGCs change as a function of retinal location (Figure 1E), converging toward the ventral pole (Figure S2) as described previously (Sabbah et al., 2017). At eye opening, there are fewer dorsal-preferring ON-OFF DSGCs and more nasal-preferring ON DSGCs compared to the adult, with this latter difference potentially due to an immature OFF response of ON-OFF DSGCs at eye opening (Hilgen et al., 2017; Rosa et al., 2016). Vertical-preferring (dorsal and ventral) ON DSGCs exhibit a small but significantly lower direction selectivity index (DSI) at eye opening (DSI = 0.44 ± 0.01; mean ± 95% CI) than adulthood (DSI = 0.53 ± 0.01; mean ± 95% CI). Moreover, there are no significant differences in the proportion of DSGCs that fall within each functional group or in tuning strength between adult normal-and dark-reared mice (Figure 1F and Figure S3). These results are similar for genetically-identified subsets of ON-OFF DSGCs (Figure S2 and S4 Hb9 and Drd4 data). Our results therefore indicate that vision does not play a role in the establishment or maintenance of retinal direction selectivity maps. We hypothesize that our previous results indicating reduced clustering of preferred directions at eye opening and after dark-rearing (Bos et al., 2016) were due to under-sampling and pooling of data across the entire retina, therefore not accounting for local differences (Figure S5).

These findings indicate direction selectivity maps are well-established at eye opening, independent of visual experience. Our results are consistent with previous studies showing that, unlike in ferrets (Li et al., 2006), dark-rearing in mice does not alter the distribution of preferred directions of DSGCs in cortical layer 2/3 compared to normal-reared adults (Hagihara et al., 2015; Rochefort et al., 2011) (Note: cortical direction selectivity is partly inherited from the retina (Hillier et al., 2017)).

We next explored whether spontaneous activity plays a role in the development of direction selectivity maps. Before eye opening, the primary driver of retinal activity stems from retinal waves, which are spontaneously-produced waves of activity that propagate across the surface of the retina (Feller et al., 1996; Maccione et al., 2014). During the first postnatal week, retinal waves are mediated by cholinergic circuits and exhibit a propagation bias in the direction of forward optic flow (Ackman et al., 2012; Stafford et al., 2009). In mice where the β2 subunit of the nicotinic acetylcholine receptor is genetically-ablated (β2-nAChR-KO), retinal waves are severely disrupted (Burbridge et al., 2014; Stafford et al., 2009). Moreover, β2-nAChR-KO mice lack a horizontal optokinetic reflex (Wang et al., 2009), which is dependent on retinal direction selectivity (Shi et al., 2017; Yoshida et al., 2001). We therefore assessed direction selectivity maps in the retinas of β2-nAChR-KO mice.

At eye opening, β2-nAChR-KO mice exhibit a dramatic underrepresentation of horizontal-preferring DSGCs (Figures 2A and 2C) and a slight reduction in the tuning strength of both horizontal-and vertical-preferring DSGCs (Figure 2D). This absence of horizontal direction selectivity persists into adulthood. These results are similar when we analyze individual functional subtypes of ON-OFF and ON DSGCs (Figure S6). Vertical-preferring DSGCs still map along the axes of optic flow, though there is a small but significant increase in the magnitude of skewing along the vertical axes compared to the wild type mice (Δ ∼ 3-5 °/mm; Figure 2B).

**Figure 2.**
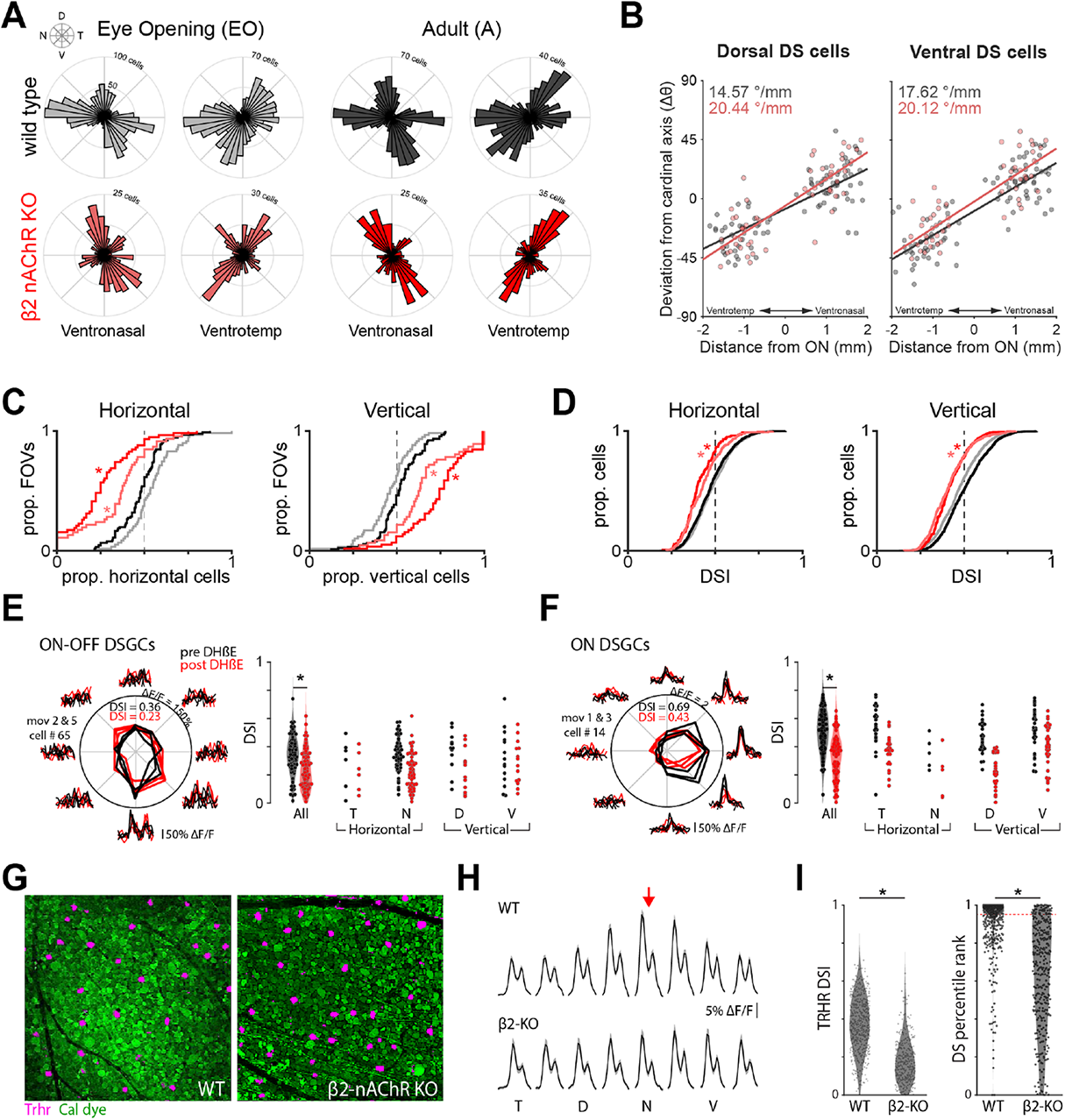
Retinal waves set up the horizontal component of the direction selectivity map. **A**. Summary data presented in polar histograms depicting the DS maps of wild type (gray/black) and β2-nAChR-KO mice (pink/red) at eye opening (gray/pink) and adulthood (black/red). The maps are segregated based on retinal location (ventronasal and ventrotemporal). Here, ON-OFF and ON DSGCs are combined into one plot because the effect occurs in both subtypes (See Figure S6). N for EO β2-nAChR-KO mice = 741 DSGCs across 5 mice; N for adult β2-nAChR-KO mice = 781 DSGCs across 8 mice. **B**. Summary data showing the rate of change of a functional group’s preferred direction as a function of distance from the optic nerve for adult WT (black) and β2-nAChR-KO (red) mice. **C**. Cumulative distribution plots depicting the different proportions of horizontal-or vertical-preferring ON-OFF and ON DSGCs per field of view. * different from both EO WTs and adult WTs, p < 0.01. **D**. Cumulative distribution plots depicting the direction selectivity index (DSI) values of horizontal-or vertical-preferring ON-OFF and ON DSGCs. * different from both EO WTs and adult WTs, p < 0.01. **E/F**. Example tuning curve (left) and summary data (right) showing ON-OFF (E) and ON DS (F) tuning before and after the application of DHβE in WT adult mice. N = 104 ON-OFF DSGCs and 96 ON DSGCs across 3 mice. * p = 4*10^−4^& 3*10^−10^for main effect of drug for ON-OFFs and ONs, respectively. No significant interaction between drug and functional cell group. ANOVA. **G**. Two example field of views showing Trhr-GFP DSGCs in magenta in mice that exhibit normal waves (WT; left) and mice that exhibited reduced cholinergic waves (β2-nAChR-KO; right). **H**. Average response of Trhr-GFP DSGCs to bars moving in different directions in both the WT (top) and β2-nAChR-KO (bottom) mice. Shaded area depicts one standard deviation from the mean. Red arrow is the typical preferred direction of Trhr-GFP DSGCs (180°, nasal). **I**. Summary data for DSI (left) and the percentile rank of each cell’s DSI compared to permutations where the directions of the moving bar are block-shuffled (right). For reference, 95^th^percentile is considered statistically-significantly DS (red dashed line). N = 465 Trhr-GFP DSGCs across 2 WT mice and 338 Trhr-GFP DSGCs across 2 β2-nAChR-KO mice. * DSI: p = 2.23*10^−79^; percentile rank: p = 1.22*10^−58^, unpaired t-test.

To better understand the mechanisms that contribute to the loss of horizontal direction selectivity in β2-nAChR-KO mice, we performed the following two experiments: First, we tested for an acute role of cholinergic signaling, since it is known to contribute to the direction-selective computation (Sethuramanujam et al., 2016). Pharmacological block of the β2 subunit of the nAChR using Dihydro-β-erythroidine (50 µM DHβE) significantly reduces but does not abolish the DSI of all DSGCs, regardless of preferred direction (Figure 2E). Therefore, acute loss of cholinergic signaling does not explain the dramatic loss of horizontal direction selectivity in β2-nAChR-KO mice.

Second, we assessed the response properties of nasal-preferring DSGCs in β2-nAChR- KO mice in which a known population of nasal-preferring DSGCs express GFP under the Trhr promoter (Figure 2F). In the β2-nAChR-KO:Trhr-GFP mice, Trhr-GFP DSGCs are responsive to drifting bars moving in all directions (Figure 2G) and have significantly lower DSI values compared to the WT:Trhr-GFP mice (Figure 2H, left). Hence, in β2-nAChR-KO:Trhr-GFP mice, many Trhr-GFP DSGCs are not direction-selective (Figure 2H, right). These data indicate that in β2-nAChR-KO mice, nasal-preferring DSGCs are likely to have weaker inhibitory input for null-direction stimulation than WT mice.

The finding that early retinal activity impacts horizontal, but not vertical, direction selectivity is reminiscent of a recent study of a human disease model for nystagmus, the FRMD7-KO mouse. Indeed, in both the β2-nAChR-KO and FRMD7-KO mice, the horizontal optokinetic reflex is missing while the vertical reflex is spared (Wang et al., 2009; Yonehara et al., 2016). Together these findings indicate that different mechanisms underlie the development of horizontal vs. vertical direction selectivity. One hypothesis is that the asymmetric inhibitory circuits that mediate vertical direction selectivity mature later, potentially depending on late stage glutamatergic waves, which are spared in the β2-nAChR-KO mouse. Alternatively, the distribution of preferred directions along the vertical axis may rely on molecular gradients, like the EphB and their ephrin ligands, that are expressed transiently in the retina during the same developmental period when direction-selective circuits develop (Thakar et al., 2011).

The reduction of horizontal directional selectivity upon disruption of retinal waves is perhaps the most dramatic impact yet reported on the role of spontaneous retinal activity in retinal circuit development. These results have implications for the influence of retinal activity on the development of direction selectivity in higher visual circuits. Indeed, there is growing evidence that early spontaneous activity is critical in establishing cortical circuits (Moreno-Juan et al., 2017; Seabrook et al., 2013; Wang et al., 2021), but our findings highlight the possibility that the locus of plasticity occurs in circuits upstream of cortex.

One intriguing hypothesis is that retinal waves might provide an instructive cue to developing direction-selective circuits. Retinal waves mimic the type of optic flow that animals will eventually experience as they navigate their environment. Indeed, the optic flow experienced by Xenopus tadpoles, an animal that does not exhibit retinal waves (Demas et al., 2012), instructs their retinotopy (Hiramoto and Cline, 2014). Therefore, retinal waves confer directional information onto developing retinal circuits before conventional photoreceptors come online, and in so doing contribute to the establishment of visual maps before mice can see.

## Acknowledgements

We thank Benjamin E. Smith for technical support and members of the Feller lab for commenting on the manuscript.

## Author contributions

Conceptualization, A.T. and M.B.F.; Methodology, A.T. and M.B.F.; Software, A.T.; Formal Analysis, A.T., K.B., and M.B.F.; Investigation, A.T. and K.B.; Writing – Original Draft, A.T. and M.B.F.; Writing – Review & Editing, A.T., K.B., and M.B.F.; Visualization, A.T. and M.B.F.; Funding Acquisition, A.T. and M.B.F.;

## Data and materials availability

All custom-written software described in the data analysis section are available online: https://github.com/atiriac/Alex-DS-project.The processed dataset (∼1-2 GB) and the unprocessed dataset (>300 GB) generated during this study are already on the author’s online Dropbox server and will be shared publicly upon acceptance. Further information and requests for resources and reagents should be directed to and will be fulfilled by the Lead Contact, Marla B. Feller.

## Supplementary Materials

Materials and Methods

Figures S1-S6

## Supplementary Material

### DATA AND SOFTWARE AVAILABILITY

All custom-written software described in the data analysis section are available online: https://github.com/atiriac/Alex-DS-project

Further information and requests for resources and reagents should be directed to and will be fulfilled by the Lead Contact, Marla B. Feller.

### EXPERIMENTAL MODEL AND SUBJECT DETAILS

#### Animals

All animal procedures were approved by the UC Berkeley Institutional Animal Care and Use Committee and conformed to the NIH *Guide for the Care and Use of Laboratory Animals*, the Public Health Service Policy, and the SFN Policy on the Use of Animals in Neuroscience Research. For the majority of experiments, we used C57B6 mice. In a subset of experiments, we used transgenic mice where Hb9 cells (ventral-preferring DSGCs) or Drd4 cells (nasal-preferring DSGCs) were tagged with a GFP marker. These transgenics were on the C57B6 genetic background. To study the role of spontaneous activity, we used the β2-nAChR-KO mouse where the beta subunit of the nicotinic acetylcholine receptor is knocked out.

To study the effects of visual deprivation on the development of direction selectivity maps, mice were born and raised in rooms that either had 12-hour day/night cycle (NR and EO groups) or 24 darkness (DR group). For the dark-reared mice, all animal husbandry was conducted with red light, which minimizes stimulation of photoreceptors.

### DATA ACQUISITION DETAILS

#### Retina preparation

Mice were deeply anesthetized with isoflurane inhalation and euthanized by decapitation. Eyes were immediately enucleated and retinas were dissected in oxygenated (95% O_2_/ 5% CO_2_) Ames’ media (Sigma) at room-temperature under infra-red illumination. Cuts along the choroid fissure were made prior to isolating the retina from the retinal pigmented epithelium. These cuts were made to precisely orient retinas and reduce orientation variability between preparations (Stabio et al., 2018). Isolated retinas were mounted whole over white filter paper (Whatman) with the photoreceptor layer side down, and transferred in a recording chamber of an upright microscope for calcium dye loading and subsequent imaging. The whole-mount retinas were continuously perfused (3 ml/min) with oxygenated Ames’ media at 32-34°C for the duration of the experiment. Retinas were bolus loaded with either the green calcium dye Cal 520 AM or the red calcium dye Cal 590 AM. The retina from the other eye was kept in the dark at room temperature in Ames’ media bubbled with 95% O_2_, 5% CO_2_ until use (maximum 8 h).

#### Two-photon calcium imaging

Two-photon fluorescence measurements were obtained with a modified movable objective microscope (MOM) (Sutter instruments, Novator, CA) and made using a Nikon 16X, 0.80 NA, N16XLWD-PF objective (Nikon, Tokyo, Japan). Two-photon excitation of calcium dyes was evoked with an ultrafast pulsed laser (Chameleon Ultra II; Coherent, Santa Clara, CA) tuned to 920 nm for green dyes and GFP or 1040 nm for red dyes. The microscope system was controlled by ScanImage software (www.scanimage.org). Scan parameters were [pixels/line x lines/frame (frame rate in Hz)]: [256 x 256 (2.96Hz)], at 1 ms/line. This MOM was equipped with a through-the-objective light stimulation and two detection channels for fluorescence imaging.

A previous study reported that a 2-photon laser at appropriate power for imaging (5-10 mW) can induce light responses (Euler et al., 2009). To account for 2-photon-mediated responses, we adapt retinas to the laser with a 5-minute imaging session, a strategy outlined in the original paper.

To keep track of the location of the retina, we zeroed the micromanipulator that moves the objective on the optic nerve, and oriented ventronasal retina along the x axis, and ventrotemporal along the y axis (or vice versa). For every field of view, we kept track of the x and y distance from the optic nerve. At the end of the experiment, we replaced the objective with a 10X objective and collected bright field images of the whole retina. We used this image of the whole retina to determine how radially offset ventronasal and ventrotemporal retina were from the x and y axis. Across all of our preparation, ventronasal and ventrotemporal retina were offset from the x and y axis by an average of -4.9°with a standard deviation of 11.9°.

#### Interline visual stimulation

Visual stimuli patterns were generated in matlab using the psychphysics toolbox and projected onto the retina using a digital micromirror device that contains an LED (UV: 375 nm). To decrease the background signal entering the photomultiplier tubes due to UV stimulation of the calcium dye, the stimuli was delivered on the flyback of the fast axis scanning mirror during a unidirectional scan so as to interleave the stimuli with the imaging. The rate at which the visual stimulus was shown with the interline (1 KHz) is faster than the flicker fusion frequency for mice (approximately 15-20 Hz) (Tanimoto et al., 2009). The intensity of the UV stimulus was 2×10^6^photons s^-1^um^-2^. Our stimulus was a rectangle (width = 500 µm, length = 1000 µm) that drifted across the field of view in 8 different directions (every 45 degrees) at speed of 250 um/s, which is ideal for activating both On and On-Off DSGCs (Dhande et al., 2013; Oyster, 1968). Each direction was repeated 3 times in a block shuffled manner so that all 8 directions were presented before repeating, with 10 seconds of downtime between stimuli. In all cases, the stimuli had a 100% positive contrast (bright on dark background).

The same MATLAB code that generated the visual stimuli also generated a text file that contained metadata for every trial, like the width/length/speed of the bar, the direction the bar was moving in during each trial, and the start and end time of each trial.

#### Pharmacological experiments

We blocked the α_4_β_2_ subunits of the nicotinic acetylcholine receptors via the application of DHβE (10 µM; Tocris, part 2349).

### DATA ANALYSIS DETAILS

#### Image processing

Raw movies were motion-corrected and normalized into ΔF/F_0_ automatically using a custom-made FIJI macro that was run in ImageJ v1.52n. Briefly, 1) movies were motion corrected on a duplicate of the raw data that had been averaged in the time dimension (zMean = 15 seconds). 2) Frames where the light stim occurred were removed to isolate baseline F. 3) the baseline F was subtracted from the raw F movie, and this result was divided by the baseline F. The resulting ΔF/F_0_ movies were then transferred to MATLAB for further image analysis.

#### Semi-automatic detection of DSGCs

A MATLAB code was written to automatically identify potential ROIs within a ΔF/F_0_ movie that were direction selective. Briefly, the code analyzes every pixel in the x-y dimension by 1) taking a mean average of its neighbor pixels, 2) computing the average peak ΔF/F_0_ response for every stimulus direction and 3) computing the vector sum (VS) and direction selectivity index (DSI) for each pixel.

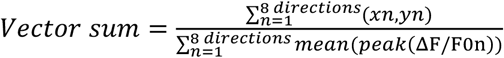

Where x_n_ and y_n_ are the cartesian coordinates of the polar vector where the direction of the vector is the stimulus direction and the length of the vector is the average peak ΔF/F_0_ for that direction. The direction of the vector sum is the ROI’s preferred direction, which is used to calculate DSI, and the length of the vector sum is the magnitude of the tuning.

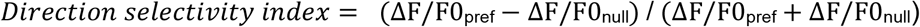

Where pref is the direction angle closest to the vector sum’s direction and null is 180°rotated from pref.

Next the MATLAB code used a 2D median filter to enrich the cell-like ROIs that exhibit similar preferred directions. The resulting image is then overlaid on an average fluorescence image of the motion-corrected movie and oval ROIs are drawn in FIJI over regions that were mathematically determined to be DS and that also correspond to an anatomical cell. These ROIs are then transferred to MATLAB for further analysis.

#### Manual classification of On-Off and On cells

A MATLAB code was written to present a user with a GUI that cycled between every identified cells. The GUI was customized to display the chronological ΔF/F_0_ trace of the cell, the 3 blocked and averaged ΔF/F_0_ traces for each of the eight different directions, and a tuning plot of the cell using the blocked traces. Using this GUI, a user classified cells as either “On-Off” if they exhibited a ΔF/F_0_ peak at the onset and offset of the moving bars, “On” if they exhibited a ΔF/F_0_ peak only at the onset of the moving bars, and “Bad” if the cells exhibited grossly inconsistent responses to each of the 3 trials for the eight different directions.

#### Statistical determination of direction selectivity

The following statistical approach was used to determine which cells were significantly direction selective: For each cell, the DSI was first calculated. Then, for 1000 permutations *in silico*, the directions of the moving bar stimuli were block-shuffled and the DSI was again calculated. The cell’s DSI calculated from the non-permuted dataset was ranked against all of the DSIs calculated from the permuted dataset. If the cell’s actual DSI ranked higher than 95% of the permuted DSIs, it was determined to be significantly direction selective.

#### Clustering analysis

We used the same clustering method that was described in a previous study (Bos et al., 2016). Briefly, K-means clustering analysis in MATLAB software was used to evaluate the pattern of distribution of the preferred directions from both On-Off and On DSGCs. Note that because the preferred directions of On DSGCs in our dataset seemed to follow the same directions as their On-Off counterparts, similar to what was described previously (Sabbah et al., 2017), we ultimately combined On-Off and On groups when performing the clustering. All the lengths of the preferred directions were fixed to 1 and these were transformed into Cartesian coordinates for subsequent angular distance measurement. This method optimizes the set of clusters with respect to the distance between each point and the centroid of its cluster, summed for all points. We compared 2-8 cluster numbers, and we calculated the fitness of clustering by using the silhouette value (SV).

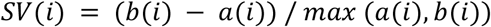

Where a(i) is the average distance between I and all other data within the same cluster (called measure of cohesion), and b(i) is the average distance between I and all points in the nearest cluster (called measure of separation from the closest other cluster). A SV close to 1 indicates data perfectly clustered, whereas a SV close to 0 reflects data which are ambiguously clustered.

Since a subset of experiments were performed in transgenic mice where known DSGCs were labelled with GFP, we used these known cell types to define the clusters. For example, the cluster that pointed ventrally and matched the Hb9-GFP, which labels a subset of ventral-preferring cells, was defined to be the ventral cluster of DSGCs. Immediately clockwise of this cluster, the cluster that pointed nasally and matched the Drd4-GFP, which labels a subset of nasal-preferring cells, was defined to be the nasal cluster. The cluster 180°rotated from the defined ventral cluster was defined as the dorsal cluster, and the cluster 180°rotated from the nasal cluster was defined as the temporal cluster. Because the preferred directions change as a function of retinal location this analysis was performed separately for ventronasal and ventrotemporal retina, as well as for central (<1000 µm from optic nerve) and peripheral fields of view (≥1000 µm from optic nerve).

#### Quantitative properties of the direction selectivity map

From the dataset that is classified as On-Off and On cells and for each directional cluster, we computed the proportion of cells, the DSI, the VS, and the variance of angles within each class of DSGCs. Cumulative distribution plots were used to present the data. Differences between groups was determined by performing an ANOVA followed by a Tukey-Kramer post-hoc test to account for unequal sample sizes.

---

**Figure S1.**
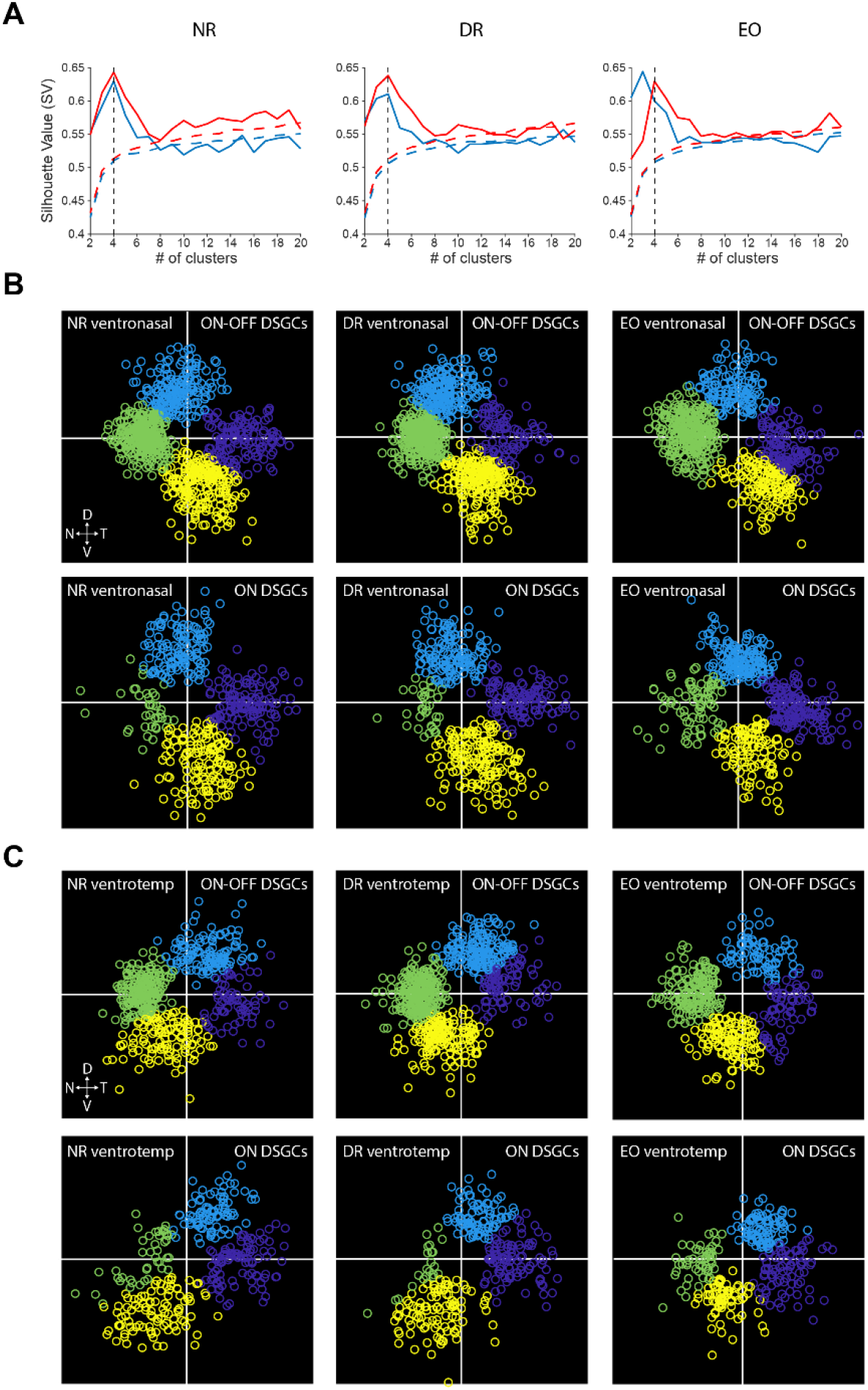
Method used to functionally cluster DSGCs. **A**. Silhouette analysis for ON-OFF (blue) and ON (red) DSGCs for the ventronasal quadrant of normal-reared (NR), dark-reared (DR), and eye opening (EO) mice. The silhouette value is a measure of how segregated the clusters are, with higher values signifying higher segregation. The red and blue dotted lines are the average of simulated data where the preferred directions were randomized. The black dotted lines indicate a cluster number of 4. **B**. Results of functionally-clustering ventronasal data into 4 groups, which are defined as temporal, nasal, dorsal, and ventral clusters. For each data point, the angle is the cell’s preferred direction, and the length from origin is its vector sum. **C**. Same as B but for ventrotemporal data.

**Figure S2.**
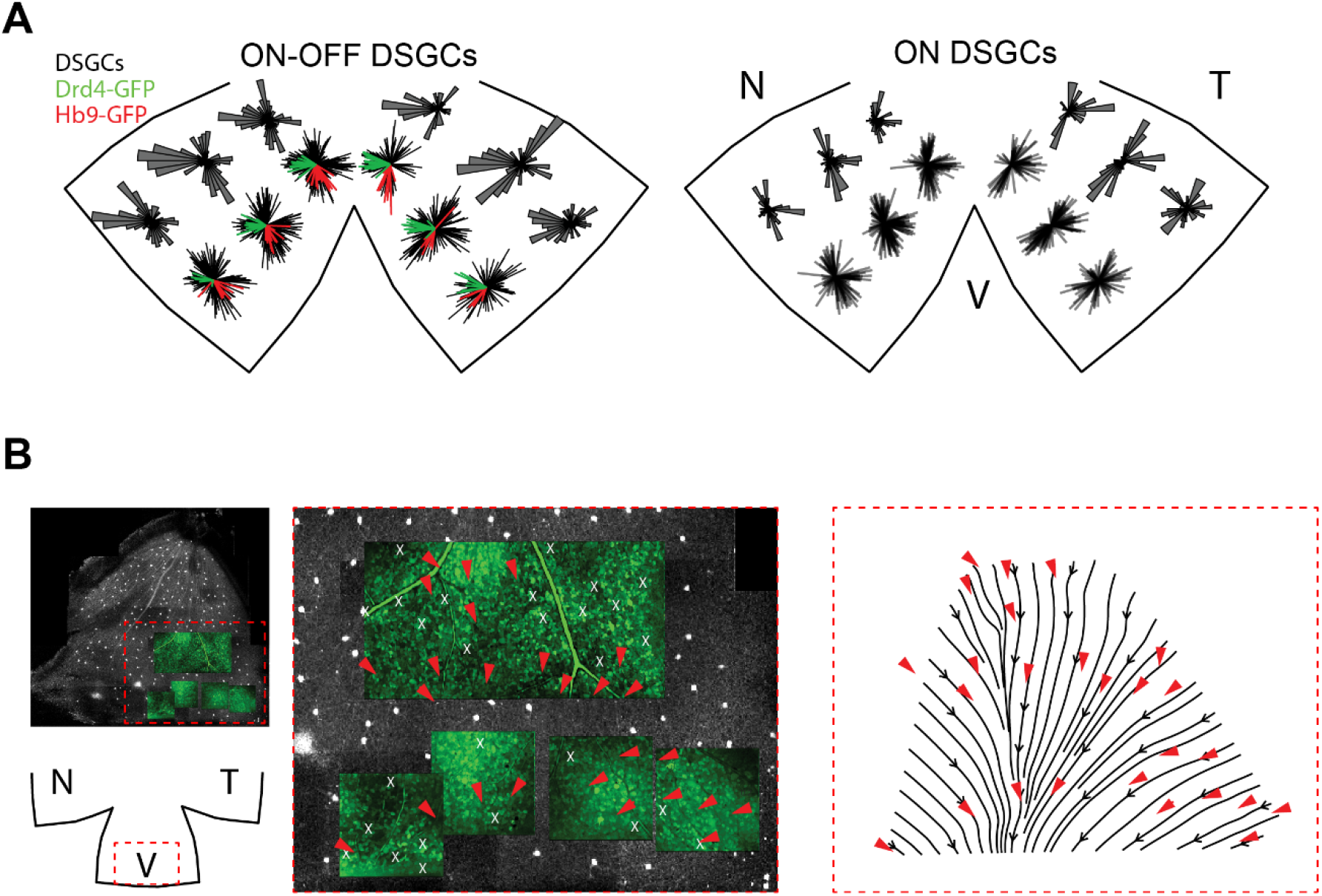
Vertical-preferring DSGCs point toward the ventral pole. **A**. Polar and rose plots depicting the local direction selectivity map at various locations of the retina. Green and red lines depict the preferred directions of genetically-identified subtypes of nasal-preferring (Drd4) and ventral-preferring (Hb9) DSGCs. **B**. Maps of the preferred direction of ventral-preferring DSGCs (Hb9-GFP) surrounding the ventral pole. Left panels depict imaging FOVs. Middle panel is a zoom in of left, with the red arrows showing the preferred directions of Hb9-GFP DSGCs. White X depict unresponsive Hb9s. Grey image in the background depicts non-imaged areas of the retina. Right panel is a vector flow map generated in MATLAB based on recorded preferred directions of the Hb9-GFP DSGCs at left. Vector flow appears to converge on ventral pole.

**Figure S3.**
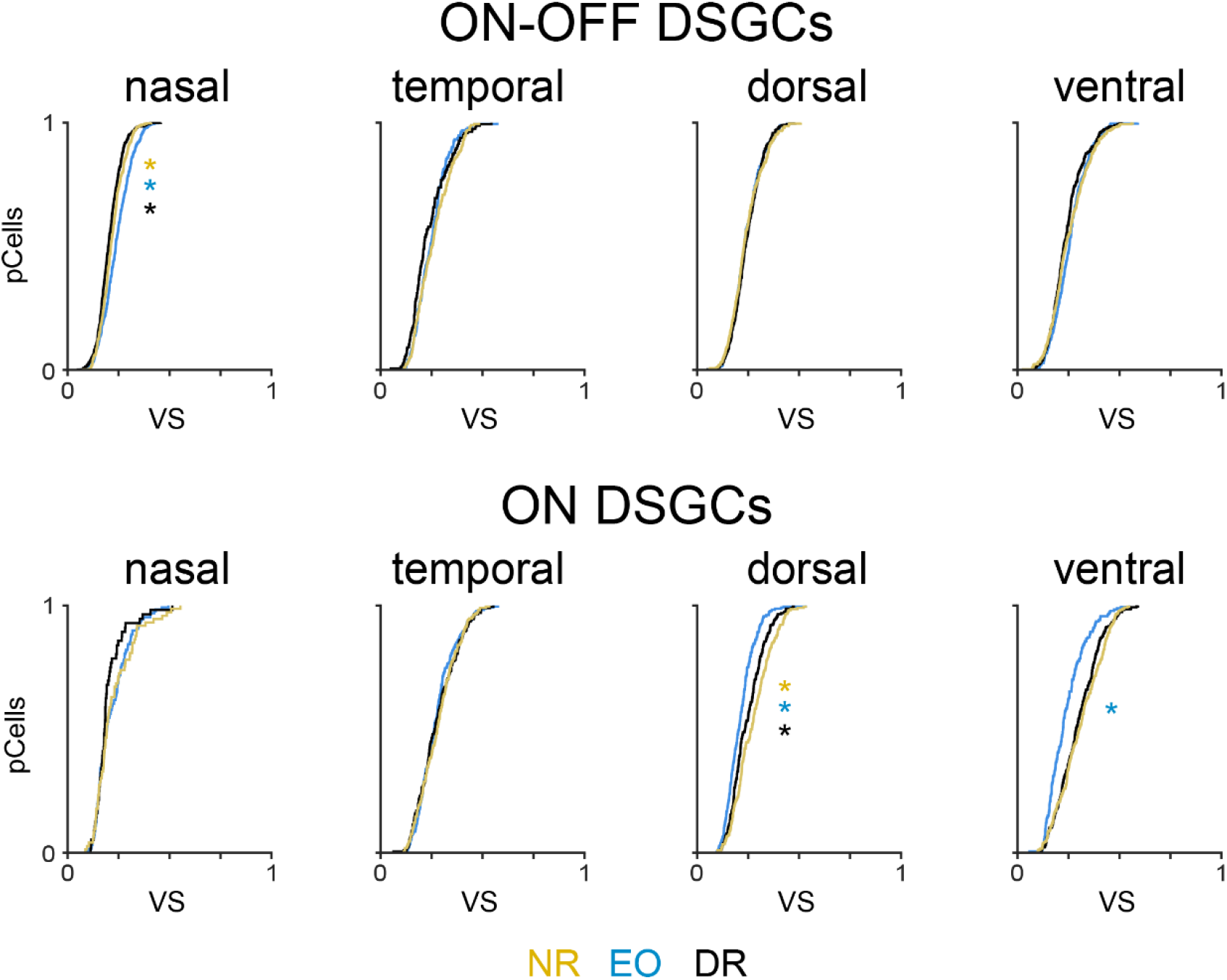
Summary results for directional tuning using vector sum. Cumulative distribution plots for vectors sum (VS) values for each DSGC subtype in retinas isolated from normally-reared mice (NR), at eye opening (EO) and dark-reared mice (DR). Vector sum is a complementary measure to direction selectivity index (DSI) of directional tuning. The significant differences in the vector sum results mirror what we observed in the DSI results. * different from other experimental group, p < 0.01.

**Figure S4.**
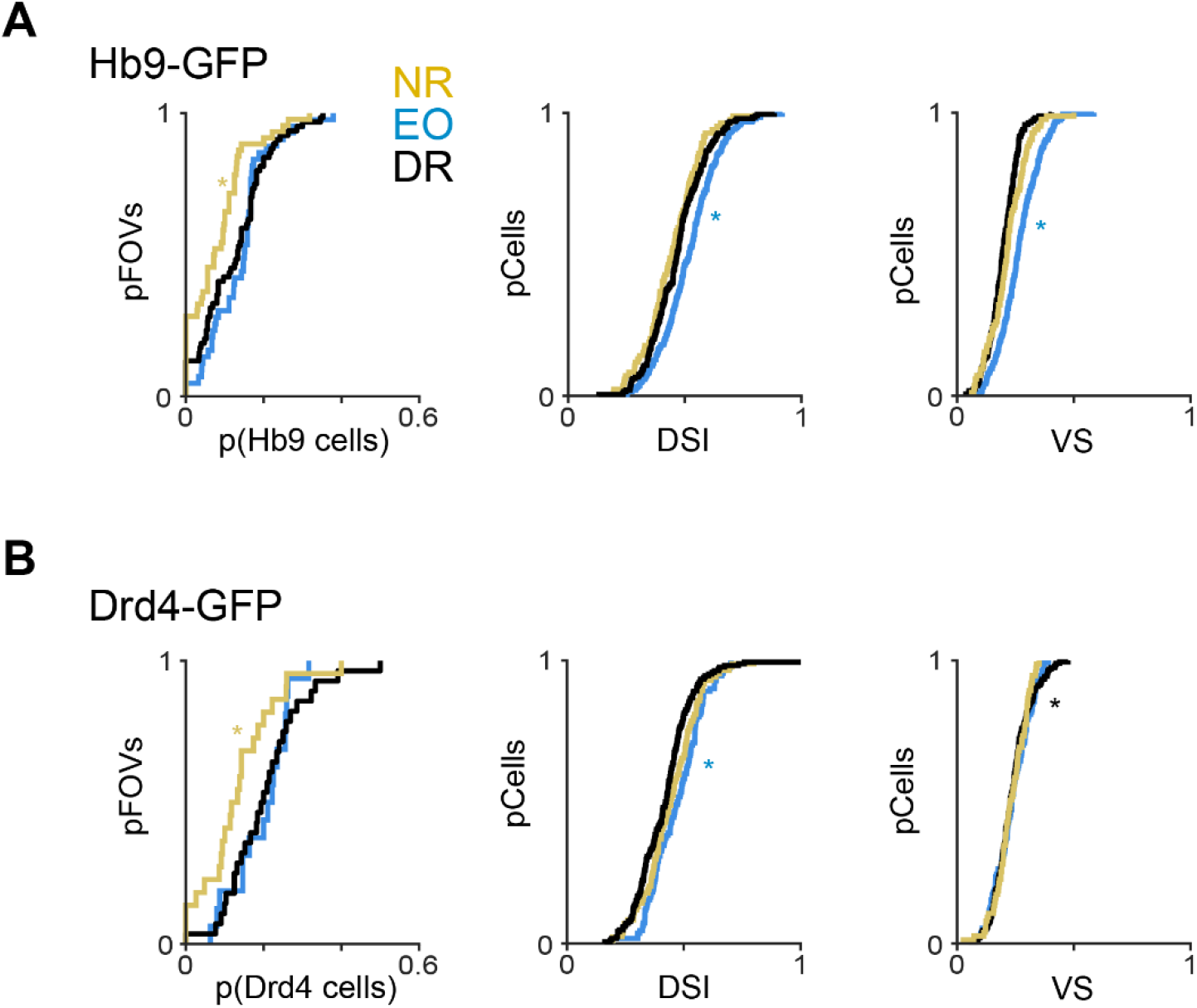
Summary results for the genetically identified subset of vDSGCs (Hb9-GFP) and nDSGCs (Drd4-GFP). **A**. Left: Cumulative distribution plots depicting the proportion of field of views that exhibit different proportions of Hb9-GFP DSGCs in retinas isolated from normally-reared mice (NR), at eye opening (EO) and dark-reared mice (DR). Middle and Right: Cumulative distribution plots for direction selectivity tuning computed as both direction selectivity index (DSI, middle) and vector sum (VS, right). FOV, field of view. * different from other experimental group, p < 0.01. **B**. Same as A but for Drd4-GFP DSGCs. In contrast to total population of DSGCs (Figure 1F), we observe a slightly but significantly lower proportion Hb9-GFP and Drd4-GFP DSGCs in normally-reared adult mice. With respect to tuning strength, Hb9-GFP and Drd4-GFP DSGCs are slightly but significantly more tuned at eye opening than in either adult group. There are no differences in tuning strength between normal- and dark-reared adults.

**Figure S5.**
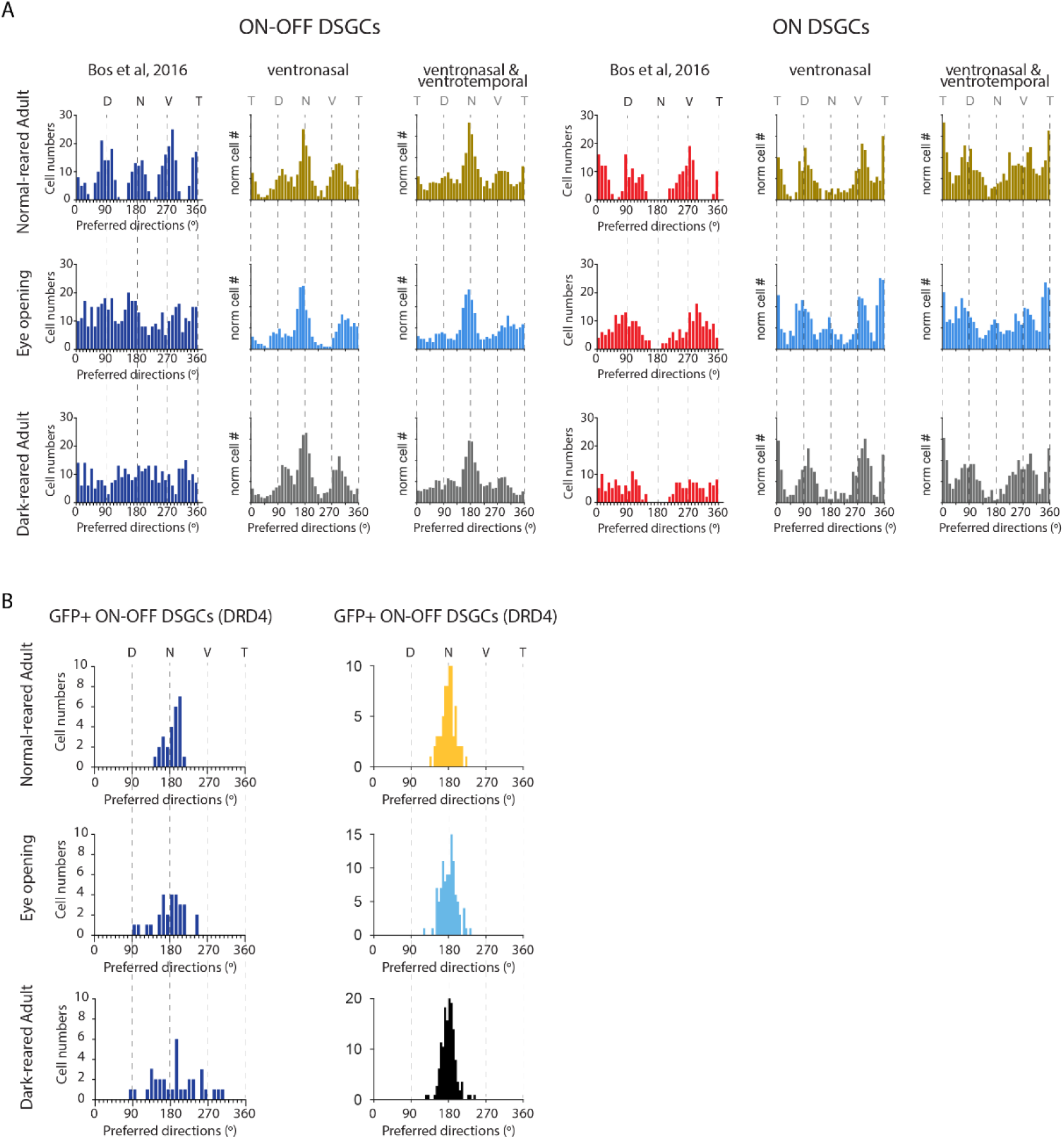
Direct comparison of distributions of preferred directions of DSGCs of this study with previous study. **A**. Left: Distribution of preferred directions from Bos et al 2016 in which we did not keep track of nasal vs. temporal location of imaging field of views (FOVs). Middle: Data from current study for FOVs in ventronasal quadrant. Right: Data from current study in which data from ventronasal and ventrotemporal are combined. Data are shown for both ON-OFF and ON DSGCs in normally-reared mice (top), at eye opening (middle) and dark-reared mice (bottom). **B**. Same as A for ON-DSGCs **C**. Distribution of preferred directions of subset of nasal-preferring DSGCs (Drd4-GFP) from Bos et al 2016 (left) and this current study (middle). Discussion: In (**Bos et al, Current Biology 2016)**, we reported that ON and ON-OFF DSGCs were directionally tuned at eye opening in mice, but that their preferred directions are not clustered along the cardinal axes as in adults. We also reported that this diffuse clustering persisted in dark-reared animals. We concluded that this data indicated that visual experience is critical for establishment of direction selectivity maps in the retina. The following year, it was demonstrated that DSGCs do not cluster along these cardinal axes as defined by motion in visual space, but rather the preferred directions of DSGCs cluster along axes defined apparent motion due to optic flow (Sabbah et al, 2017) Therefore, the distribution of preferred directions depends on the location on the retina. This direct comparison of the data from the two studies indicate that the source of the errors in the previous study are due to combining preferred directions acquired from FOVs that ranged across nasal and temporal retina and undersampling. Indeed, the use of novel calcium dyes (Cal-590 and Cal520) in this current study has greatly improved the sampling of DSGCs with significantly more DSGCs per FOV than in the previous study. This greatly improved data set indicates that the direction selectivity map is well established at eye opening and not impacted by visual experience.

**Figure S6.**
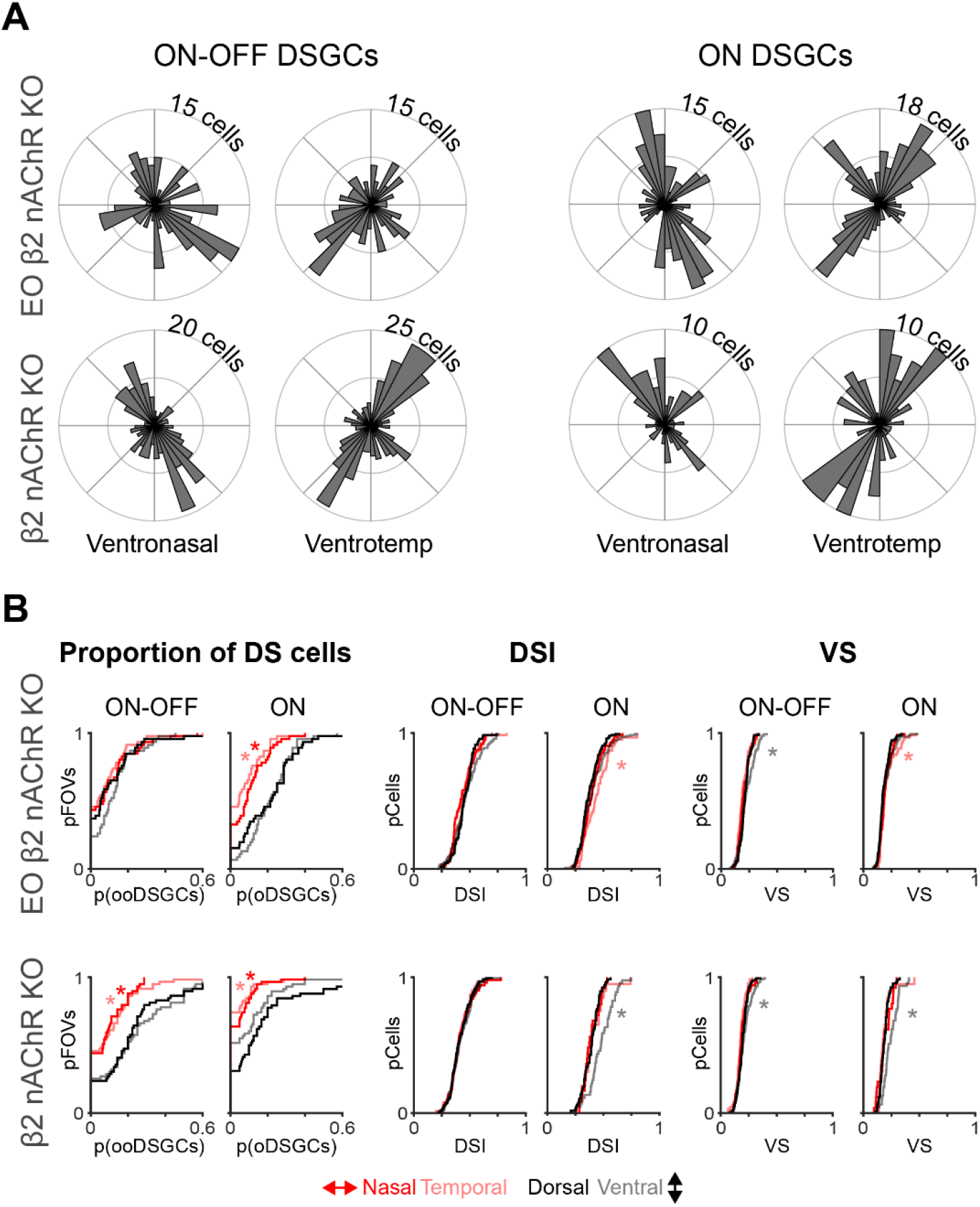
β2-nAChR-KO lack horizontal DSGCs in both ON and ON-OFF DSGCs. **A**. Polar histograms depicting the DS maps of β2-nAChR-KO mice at eye opening (top) and adulthood (bottom) for ON-OFF and ON DSGCs in ventronasal and ventrotemporal retina. **B**. Left: The proportion of field of views that exhibit different proportions of functional subtypes for ON-OFF and ON DSGCs of β2-nAChR-KO mice at eye opening (top) and adulthood (bottom). Middle & Right: same as left but for direction selectivity index (DSI) and vector sum (VS). * different from other experimental group, p < 0.01.

